# A dual-clam species 63K SNP array for sustainable production and conservation of wild resources

**DOI:** 10.64898/2026.02.06.704329

**Authors:** Marialaura Gallo, Massimiliano Babbucci, Sergio Fernández Boo, Tim Bean, Giulia Dalla Rovere, Morgan Smits, Carolina Peñaloza, Ross Houston, Shernae Woolley, Francesco Cicala, Rafaella Franch, Serena Ferraresso, Ilaria Nai, Tomaso Patarnello, Adesuyi Emmanuel Omole, Andres Blanco, Ines Sambade, Paulino Martinez, Luca Bargelloni, Massimo Milan, Luca Peruzza

## Abstract

Bivalves play an essential role in coastal ecosystems and their aquaculture represents an important economic sector in Europe playing a pivotal role within the EU Blue Growth Strategy. Among clam species, the Manila clam, *Ruditapes philippinarum*, and the grooved carpet shell, *R. decussatus*, are within the top five species in terms of production volume and economic value. In this study we designed and validated the first medium-density 63K single nucleotide polymorphism (SNP) array for these two commercially important species. By leveraging a new chromosome-level genome assembly for *R. philippinarum* and that of *R. decussatus*, we identified over 300 million SNPs through whole-genome resequencing and genotyping-by-sequencing strategies. After stringent filtering, we selected 49,392 high-quality SNPs for *R. philippinarum* and 14,193 for *R. decussatus* to construct a dual-species array. Array validation was carried out by genotyping 384 individuals across multiple wild populations and hatchery samples, demonstrating excellent performance, with 67.7% and 67.5% of SNPs classified as high-quality polymorphic markers for *R. philippinarum* and *R. decussatus*, respectively. Minor allele frequency, missing data rate, and inter-marker distance met stringent quality thresholds, confirming the array robustness for clam population genetics. Parentage analysis in *R. philippinarum* families highlighted significant power for pedigree reconstruction in breeding programs. This publicly available genomic resource provides a reliable, cost-effective genotyping platform to enable population genomics and advanced selective breeding, genome-wide association studies, and genetic monitoring, ultimately strengthening management of genetic diversity and sustainable farming for two key clam species.

## INTRODUCTION

Bivalves represent one of the most sustainable sources of animal protein, supporting human nutrition and local economies while maintaining a low environmental impact. Compared to land-based animal farming, aquaculture generally shows higher energy-conversion efficiency and makes use of extensive marine areas suitable for production (Gentry et al., 2017). In contrast to wild-capture fisheries, which haves a limited scope for expansion, aquaculture has globally seen a constant increase and has recently surpassed wild-capture fisheries in terms of tons landed (Commission et al., 2025). Within this context, production of extractive species, including bivalves, has more than doubled in volume since 2000 (Naylor et al., 2021). Bivalves are particularly valuable due to their high-quality, lean protein content and their richness in essential micronutrients like vitamin B12, zinc, iron, selenium, iodine, and omega-3s. As filter feeders, they also provide important ecosystem services by removing excess nutrients from coastal waters and thus preventing eutrophication (Gentry et al., 2017).

In Europe, bivalve aquaculture represents a major economic sector. Notably, two closely-related species, the grooved carpet shell (*R. decussatus)* and the Manila clam (*R. philippinarum*), accounted for approximately 25% of shellfish production in 2022, highlighting their economic relevance within European coastal aquaculture (Commission et al., 2025). *R. decussatus* is native to the Mediterranean Sea and Atlantic coasts of North Africa and Europe, whereas *R. philippinarum* originated from the Indo-Pacific region (Anacleto et al., 2014) and was introduced in Europe in the 70s (Delgado & Perez-Camacho, 2007) due to its faster growth and higher eurythermal abilities compared to the native species (Anacleto et al., 2014). Since its introduction, aquaculture of *R. philippinarum* has rapidly expanded from the Mediterranean to Atlantic coasts (Chiesa et al., 2017; Gosling, 2008), with Italy as the leading European producer. Today it is the most commercially valuable clam species in Europe, playing a central role in coastal aquaculture systems (Astorga, 2014).

Over the last decades, however, both species have experienced pronounced population declines across much of their distribution ranges. In the case of *R. philippinarum*, the availability of natural seed has decreased sharply in several major farming areas, largely as a consequence of climate-driven environmental stress, habitat degradation, overexploitation and increasing predation pressure from invasive species such as the blue crab (Garrabou et al., 2009; Munari, 2011; Oliver et al., 2018; Smith et al., 2021). For instance, in the Venice lagoon alone, annual production dropped from approximately 40,000 tons in the early 2000s to less than 3,000 tons in 2019. Similarly, *R. decussatus* populations have undergone severe contractions as a result of habitat loss, climate change and competition with the non-indigenous *R. philippinarum* (Chiesa et al., 2017; Ruano et al., 2015), leading to an estimated 70% reduction in landings since their peak in the early 1990s (FAO, 2024). At present, commercial production of *R. decussatus* is largely restricted to a few areas of Portugal and Tunisia.

Beyond their social and economic impacts, this decline raises serious concerns regarding the conservation and management of genetic resources. Reductions in population size, habitat fragmentation and hatchery-based seed production may lead to genetic bottlenecks, loss of local adaptation and elevated levels of inbreeding (He et al., 2025). Managing genetic resources in marine bivalves is particularly challenging due to their life-history traits (Curole & Hedgecock, 2007). Early genetic studies in bivalve populations highlighted unusually high polymorphism (S. Wang et al., 2017; Zhang et al., 2012) and frequent deviations from Hardy-Weinberg equilibrium due to high prevalence of null alleles, ultimately affecting genotyping accuracy (Hollenbeck & Johnston, 2018). Taken together, these biological features complicate assessment of population structure, genetic diversity and inbreeding, particularly in exploited or cultured populations. Moreover, disentangling natural patterns of genetic variation from anthropogenic influences, such as translocations, stock enhancement and species introductions, remains difficult in the absence of standardized, high-resolution genomic tools.

In line with the recent European Blue Growth strategy, which promotes sustainable aquaculture practices, selective breeding has emerged as a key approach to improve economically relevant traits such as growth, disease resistance, and tolerance to environmental stressors (Yáñez et al., 2022). While already widely applied in terrestrial livestock, its adoption in aquaculture is rapidly increasing, with notable successes in a few finfish species such as Atlantic salmon where selective breeding for resistance to infectious pancreatic necrosis virus (IPNV) nearly eliminated disease outbreaks within five years (Houston et al., 2020). Genetic gains in aquatic species can be higher than in terrestrial species (Yáñez et al., 2022) due to greater within-species genetic diversity (Houston et al., 2020), highlighting the untapped potential of selective breeding in aquatic species. Selective breeding in aquaculture can be implemented through a range of approaches, from traditional phenotype-based selection to advanced genomic methods such as marker-assisted and genomic selection (Houston et al., 2020). While pedigree-based and genomic approaches can substantially increase genetic gain and reduce inbreeding, their effective application critically depends on the availability of reliable genomic resources, ideally including a high-quality, contiguous reference genome and efficient genotyping tools. Genome-wide SNP panels are currently the most common tool for routine genomic evaluation (Yáñez et al., 2022) and are preferred over other types of genetic markers (Nascimento-Schulze et al., 2023), offering high accuracy, scalability and cost efficiency compared to sequencing-based approaches.

Recently, several SNP arrays have been developed for aquaculture species (Yáñez et al., 2022). In bivalves, genotyping chips are available for oysters and mussels (Gutierrez et al., 2017; Nascimento-Schulze et al., 2023), and more recently for the hard clam (*Mercenaria mercenaria*) (Grouzdev et al., 2024), a commercially important species in North America. However, no genotyping array is available for clam species of high economic relevance for European aquaculture.

To address this gap, we designed and validated a new dual-species medium-density SNP array (63K) for the Manila clam (49K SNPs) and the grooved carpet shell (14K SNPs). The array was designed using a newly obtained chromosome-level genome assembly for the Manila clam (unpublished data) and the current version of the grooved carpet shell genome (Sambade et al., 2025). Single nucleotide variants were identified through whole-genome resequencing for *R. philippinarum* and genotyping-by-sequencing (GBS) for *R. decussatus* and subsequently validated by genotyping 384 individuals (288 Manila clams and 96 grooved carpet shell) from wild European populations and hatchery samples.

This SNP array provides a reliable and cost-effective tool for high-throughput genotyping in clams, enabling accurate pedigree reconstruction in breeding programs. Furthermore, it supports population genetics applications aimed at monitoring genetic diversity and inbreeding in cultured and/or wild populations, making it feasible to disentangle the genetic basis of production and adaptive traits. By integrating this genomic resource with conventional aquaculture practices, the productivity and long-term sustainability of clam farming can be enhanced, supporting the development of more accurate breeding programs and facilitating the sustainable management of shellfish beds in both species.

## MATERIALS AND METHODS

### Sample collection and sequencing

All experimental protocols were tailored to each of the two species according to the available SNP resources and genomic tools. In both species, SNP datasets combined previously validated markers with newly generated whole-genome sequencing (WGS) data.

For Manilla clam, *R. philippinarum, i)* we used a list of 3,258 validated SNPs from a previous multispecies shellfish Affymetrix array, developed at the University of Santiago de Compostela in collaboration with Xenetica Fontao SL and Thermo Fisher Scientific, and *ii)* we performed whole-genome resequencing on individual samples from wild populations representing the major Atlantic and Mediterranean production areas in Europe (Figure 1), along with hatchery populations. Each population comprised 12–30 individuals. Total DNA was extracted from gill or muscle tissue using Invisorb® Spin Tissue Mini Kit (Invitek Molecular GmbH, Berlin, Germany) according to the manufacturer’s instructions. We generated a new chromosome-level genome assembly for this species (unpublished data), that was used as reference to align reads from whole-genome sequencing for SNP discovery and mapping, as available genomes were not resolved to chromosome level for this species. Whole genome sequencing libraries were constructed using the Illumina DNA Prep Tagmentation Kit starting from a total of 100 ng/sample. Library quantity and quality were checked with HS dsDNA QUBIT Assay and Agilent Bioanalyzer 2100 High Sensitivity DNA kit. Individual genomic DNA libraries were evenly pooled based on their concentration and were barcoded in three different pools. Pools were sequenced at the Norwegian Sequencing Centre (NSC, Norway) with Illumina NovaSeq S4 technology with 150 bp PE approach.

**Figure 1:**
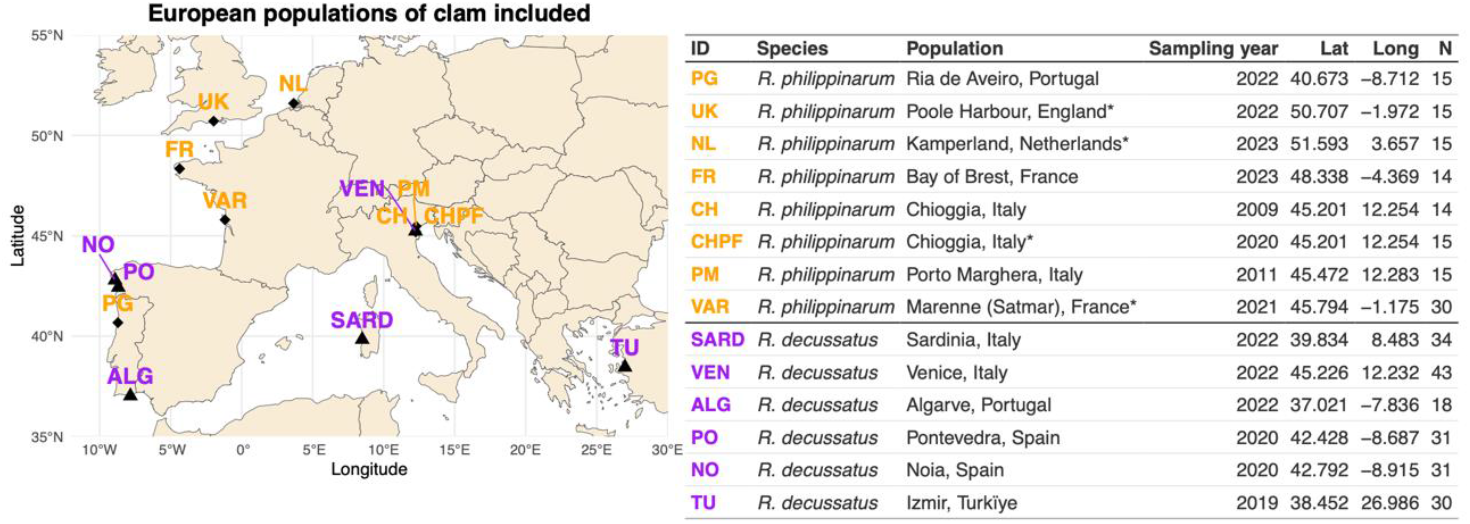
Populations included in the SNP discovery phase. Map of Europe with the names of the various populations used to design the array, colour coded by species. Details for each population (i.e. species, geographic location including latitude and longitude, sampling year and number of sampled animals) are reported in the adjacent table. “*” indicates hatchery-derived animals.

For the grooved carpet shell *R. decussatus*, two sources of SNPs were used (FigureFigure1): *i)* a validated list of 13,438 2bRAD-derived SNPs, mapped to the chromosome-level reference genome (Accession number: GCA_977926895.1), obtained from a previous population genomics study (Sambade et al., 2025) involving 215 individuals sampled from six shellfish beds roughly covering its European distribution range; and *ii)* a list of 12,813 SNPs identified from two WGS pools of 20 clams (10 males and 10 females), sampled from Muros-Noia estuary (NW Spain). For these individuals, DNA was extracted from gonad tissue using the E.Z.N.A.® DNA kit (Omega Biotek, Norcross, GA, USA) following the manufacturer’s instructions. DNA quantity and purity were estimated using a NanoDrop spectrophotometer and Agilent Bioanalyzer 2100 High Sensitivity DNA kit. WGS was performed on male and female barcoded pools at the Novogene platform (Cambridge, UK). Raw data were filtered with Trimmomatic (Bolger et al., 2014) to remove adapter sequences from library construction and perform quality filtering. Filtered reads were aligned to the reference genome using Bowtie2 (Langmead & Salzberg, 2012). SAM files were converted to BAM format using Samtools (H. Li et al., 2009). SNP were identified using the “BCFtools mpileup” pipeline. A cumulative list of ∼8 million SNPs was retrieved from both pools and subsequently filtered using custom Perl scripts. SNPs were retained if supported by a minimum read depth of 20 and with a Minor Allele Frequency (MAF) > 0.03. The SNP with the highest MAF was then selected within each 50 kb adjacent window across the whole genome, provided there was no other nearby SNP (±75bp) with MAF > 0.05.

Raw reads were filtered by quality using FastQC software v0.11.5 (http://www.bioinformatics.babraham.ac.uk/projects/fastqc/) and subsequently trimmed with FastP v1.0 (Chen, 2023) to remove adapter sequences and low-quality bases. Default parameters were applied, except for minimum read length (70 bp) and quality threshold (Q ≥ 25); the *nextera* option was applied for adapter removal. High-quality reads were aligned to the *R. philippinarum* reference genome using BWA software version 0.7.17-r1188 (Heng Li & Durbin, 2009), with the -c 1 and -M parameters enabled to ensure compatibility with downstream analyses. The SAM files produced were converted and sorted into a BAM file using SAMtools (H. Li et al., 2009). Duplicate reads were identified and removed with PicardTools v3.0.0 (http://broadinstitute.github.io/picard) using default parameters. SNP calling was performed using BCFtools v1.17 (H. Li et al., 2009) in multi-sample mode. The initial variant set was subsequently filtered with VCFtools v0.1.16 (Danecek et al., 2011) to retain only high-confidence biallelic SNPs suitable for downstream analyses.

### SNP selection for array design

#### Manila clam

For the design of the Manila clam SNP panel, we implemented a two-stage SNP selection strategy. A preliminary evaluation of the variant dataset showed that applying a mapping-quality filter (MAPQ > 20) resulted in heterogeneous depth coverage across the genome. To solve this issue, the first stage of SNP selection focused on identifying reliably characterized genomic regions (i.e., high-confidence regions), followed by a second stage in which a conventional SNP selection strategy was followed.

First, high-confidence regions were delineated using the VCF-file and consisted of segments within the assembly containing variants genotyped in at least 90% of the individuals. For the second stage, these high-confidence regions were screened for variants and subjected to QC filtering steps to generate a pool of high-quality markers used for downstream SNP selection. Variants were filtered out from the initial set of SNPs if they met any of the following criteria: (i) < 20 bp distance from the closest upstream and downstream variants, (ii) MAF < 0.05, (iii) all samples heterozygous for the given SNP, or (iv) average sequencing depth > 100 reads. MAF filtering was applied to prioritize robust and informative markers for population genetic analyses, parentage assignment and breeding applications, rather than the detection of rare variants. The flanking sequences of each candidate SNP (±35 bp) were then retrieved from the reference genome and submitted as 71-mer sequences for *in silico* probe scoring by ThermoFisher.

From the scored list of candidate markers, SNPs with at least one probeset classified as either ‘recommended’ or ‘neutral’ were retained. SNP selection was further optimized using custom scripts to prioritize markers within 20 kb windows based on informativeness and proximity to the window midpoint. This pre-selected Manila clam marker set (n=48,596) was then combined with a subset of high-quality markers from a previous Axiom array (n=796), resulting in an optimized panel with cross-platform compatibility.

#### Grooved carpet shell clam

Two quality-filtered marker datasets discovered using GBS approaches (n=13,438) and whole-genome sequencing (n=12,813), were used for SNP selection. As a subsequent filter, flanking regions of the combined dataset markers were extracted, and markers were retained if they mapped to a unique position of the chromosome-level assembly. The candidate markers were then submitted as 71-mers to ThermoFisher for *in silico* probe scoring.

The final SNP panel was assembled by prioritizing marker probesets scored as ‘recommended’, followed by ‘neutral’ ones, while requiring a minimum inter-marker distance of 10 kb between consecutive selected markers.

### SNP array validation

To validate the SNP array, a test plate with genomic DNA from 384 samples (288 from Manila clam and 96 from grooved carpet shell clam) was shipped to IdentiGEN (MSD Animal Health, Dublin, Ireland), for genotyping. For the Manila clam, DNA samples included five individuals from each of the wild populations used during the array design stage (i.e. UK, CHPF, PG, and FR), as well as several individuals from the SATMAR hatchery derived from a single mass-spawning event. For the grooved carpet shell clam, a total of 96 samples were selected from three of the beds used for SNP selection, namely Algarve (n= 18 individuals), Sardinia (n=34 individuals) and Venice (n=43 individuals). Raw data (CEL files) were processed using the Axiom Analysis Suite 5.0 software (Affymetrix) for genotype calling and quality control assessment. Samples exhibiting dish quality control (DQC) value > 0.82 and a QC call rate > 0.95, according to the *Affymetrix Best Practices Workflow*, were retained for downstream analyses. Calculations of MAF, the proportion of missing data per SNP and per individual, and inter-SNP physical distances were performed using PLINK v1.9 (Purcell et al., 2007) and GraphPad Prism 10 (www.graphpad.com). Principal component analysis (PCA) was conducted in R using the dartR package (Mijangos et al., 2022)

Parental assignment for *R. philippinarum* was carried out using the Family Assignment Program (FAP) (Taggart, 2006), which was adapted with a custom in-house script to enable the inclusion of all SNPs present in the array.

## RESULTS AND DISCUSSION

### SNP selection and array construction

For the Manila clam, the average number of 150 bp PE Illumina reads per sample was 52 million, of which 50.8 million high-quality reads were retained after trimming, with a range of 38 to 72 million raw reads per sample. Overall, 94% of the retained reads mapped to the reference genome. SNP calling via BCFtools identified a total of 330,471,516 biallelic SNPs (MAPQ score > 20) as candidates for the array. These candidate markers were evaluated using a two-stage SNP selection process. In the first stage, a total of 1,302,103 high-confidence genomic regions were identified across the Manila clam genome. In the second stage variant screening within these regions yielded 824,087 high-quality *de novo* markers, which together with the 3,258 validated SNPs from a previous array, were submitted to Thermo Fisher Scientific for *in silico* evaluation. For the grooved carpet shell, the initial lists of 13,438 RAD-derived SNPs and 21,086 WGS SNPs were filtered to retain only probes uniquely aligning to the reference genome, which were sent to Thermo Fisher for *in silico* evaluation. The final SNP array comprised 63,585 markers, including 49,392 for *R. philippinarum* and 14,193 for *R. decussatus*, evenly distributed across the chromosomes (Figure 2).

**Figure 2:**
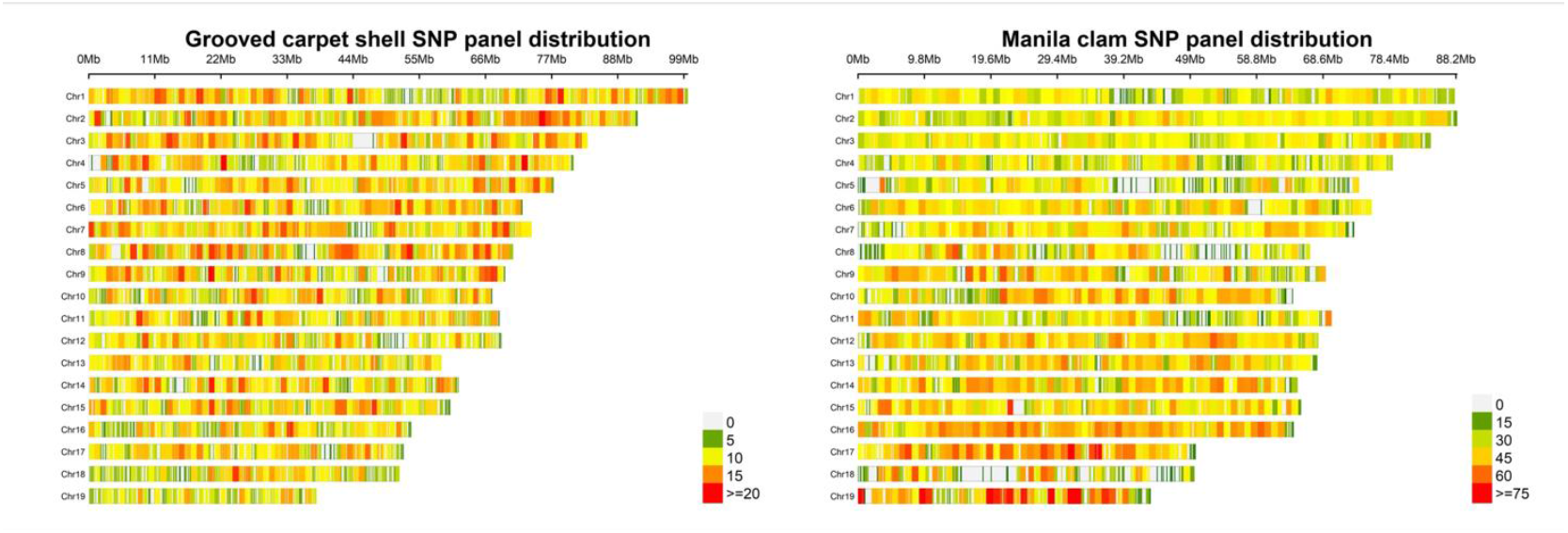
Probe distribution across chromosomes for each species (heatmap).

Functional annotation using Annovar (K. Wang et al., 2010) revealed differences in marker distribution between the two species, mainly in exonic (26.2% vs 12.0% for *R. philippinarum* and *R. decussatus* respectively), and intronic (25.6% vs 38.4%) regions. Minor differences were observed in intergenic regions (38.1% vs 39.7%), and in less represented categories such as 3’UTR, 5’UTR, and other genomic regions (Figure 3). Observed differences in the distribution of SNPs might reflect disparities in genome annotation completeness, divergent genomic architecture, or the distribution of AlfI restriction sites, the main source of SNPs for *R. decussatus* from 2b-RADseq (Sambade et al., 2025). Although the SNP array contains more markers for *R. philippinarum*, it is expected to have broad applicability for pedigree reconstruction, genome-wide association studies and genomic prediction of economically important traits in breeding programs and for population genomics studies and adaptation in both species. This enables investigations into the genetic architecture of complex traits, including resistance to marine pathogens and tolerance to environmental stressors (e.g. elevated temperatures and salinity fluctuations), as well as applications in population genetics and conservation studies.

**Figure 3:**
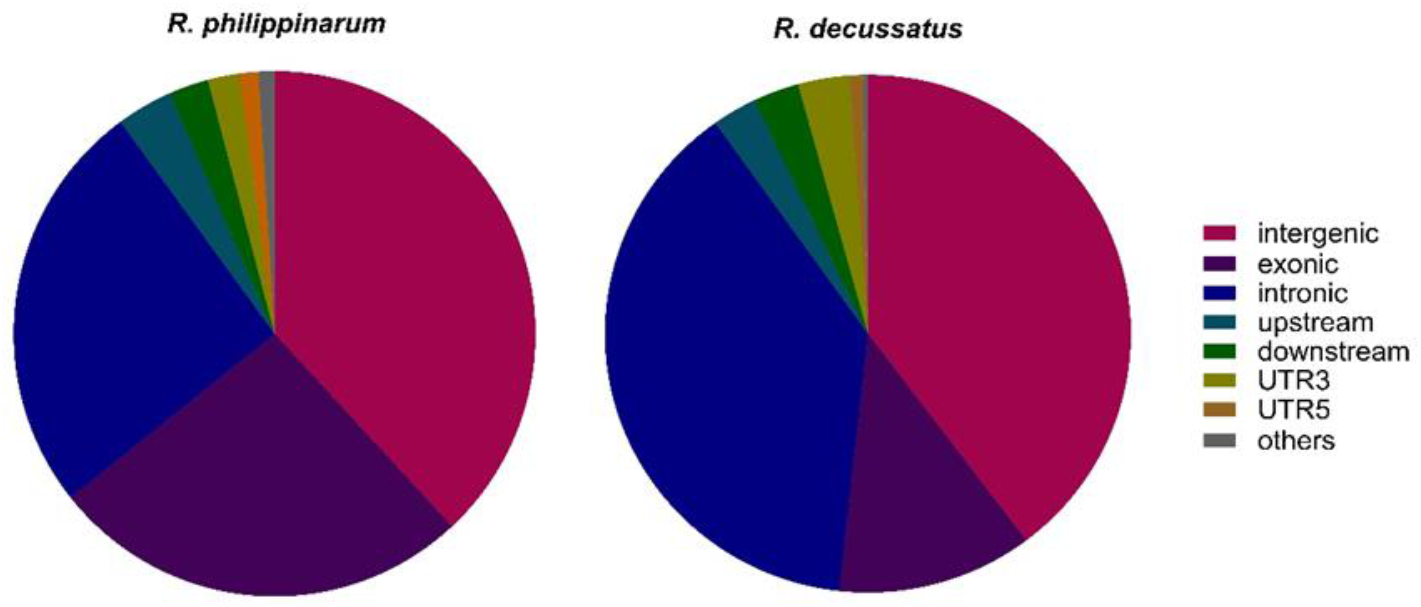
Marker composition of the 63k SNP clam array

### Evaluation of SNP array performance

To assess the performance of the multi-species SNP array, a total of 288 *R. philippinarum* and 96 *R. decussatus* clams from different locations across the European distribution range of both species were genotyped (Figure 1). Quality control filtering showed high sample retention rate, with 276 out of 288 *R. philippinarum* (95.8%) and 95 out of 96 *R. decussatus* (98.9%) samples passing Axiom QC metrics. Using the Axiom Best Practices Workflow, SNPs were assigned to six categories: *PolyHighResolution* (polymorphic SNPs with clear genotype separation), *MonoHighResolution* (monomorphic SNPs), *CallRateBelowThreshold* (SNPs with call rates <97%), *NoMinorHom* (polymorphic SNPs lacking one homozygous class), *OffTargetVariant* (OTV; displaying atypical genotype clustering pattern), and *Other* (SNPs with multiple clustering issues). For *R. philippinarum*, 31,121 polymorphic sites (63%), 2,136 sites with no minor allele homozygotes (4.3%), and 196 monomorphic sites were identified (0.4%), yielding 33,453 high-quality markers (67.7%) (Table 1). For *R. decussatus*, 8,622 polymorphic sites (60.7%), 814 sites with no minor allele homozygotes (5.8%), and 151 monomorphic sites (1.1%) were retained, yielding a total of 9,587 high-quality and recommended markers (67.54%) (Table 1). The large number of high quality markers (i.e. “Poly High Resolution” and “No Minor Hom” markers) is in line with previous SNP arrays developed for hard clam (*Mercenaria mercenaria*) (Grouzdev et al., 2024), the Eastern oyster (*Crassostrea virginica*) (Xuereb et al., 2023), the Pacific oyster (*Magallana gigas*) (Qi et al., 2017) and the European oyster (*Ostrea edulis*) (Gutierrez et al., 2017).

**Table 1.**
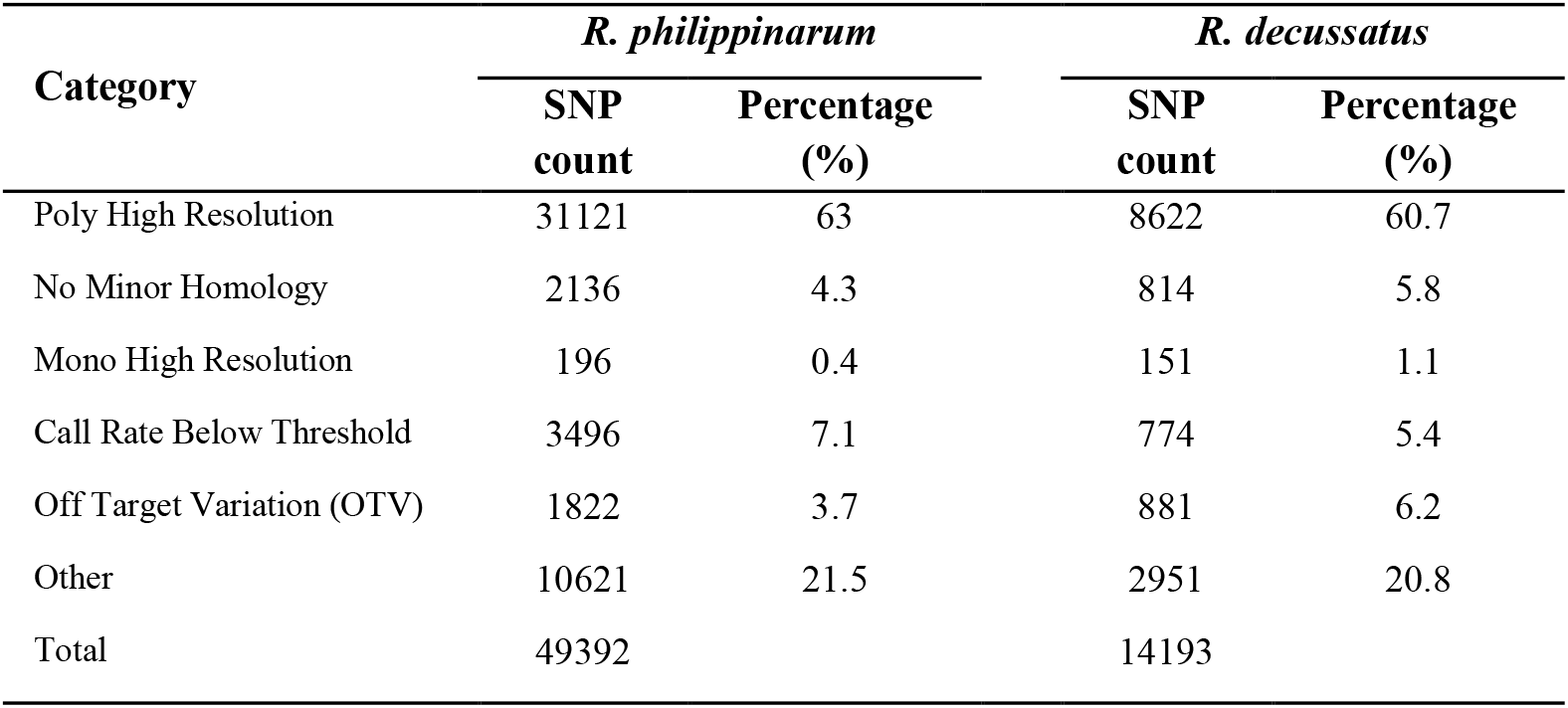
SNP quality classification in the 63K SNP Array for Manila clam and grooved carpet shell.

| Category | *R. philippinarum* |  | *R. decussatus* |  |
| --- | --- | --- | --- | --- |
|  | SNP count | Percentage (%) | SNP count | Percentage (%) |
| Poly High Resolution | 31121 | 63 | 8622 | 60.7 |
| No Minor Homology | 2136 | 4.3 | 814 | 5.8 |
| Mono High Resolution | 196 | 0.4 | 151 | 1.1 |
| Call Rate Below Threshold | 3496 | 7.1 | 774 | 5.4 |
| Off Target Variation (OTV) | 1822 | 3.7 | 881 | 6.2 |
| Other | 10621 | 21.5 | 2951 | 20.8 |
| Total | 49392 |  | 14193 |  |

Leveraging this subset of high-confidence SNPs, we conducted several performance evaluations in both species. MAF distributions were consistent with expectations following the application of a MAF ≥ 0.05 threshold: for *R. philippinarum*, only 3,138 of the 33,453 SNPs exhibited MAF below this threshold, and for *R. decussatus* the number was even lower, with only 532 out of 9,587 SNPs falling below 0.05. The vast majority of markers in both species therefore displayed MAF values between 0.05 and 0.5.

Specifically, in *R. philippinarum* 17,880 SNPs had MAF ≥ 0.2 (53.4%), while in *R. decussatus* 6,285 (65.5%) met this criterion (Figure 4A, D), reflecting a highly informative allele frequency spectrum across the retained loci. Missing genotype rates were assessed at both the SNP and individual levels, with a maximum missing data threshold of 5%. All markers and samples in both species satisfied this criterion, and no exclusions were required at this filtering step (Figure 4B, E; Supplementary Figure S1), highlighting the high genotyping reliability of the array. Marker spacing analysis showed a denser genomic distribution in *R. philippinarum*, with a median inter-SNP distance of 19 kb (Figure 4C), compared to 85 kb in *R. decussatus* (Figure 4F). This disparity reflects the larger initial panel in *R. philippinarum* and may facilitate higher-resolution genomic analyses in this species. Despite the lower density, the inter-marker spacing in *R. decussatus* remains sufficient for genome-wide applications such as population structure and diversity studies. The development of a medium-density SNP array is particularly valuable in bivalves due to their inherently high polymorphism, frequent segregation distortions, and the prevalence of null alleles, which complicate reliable genotyping using standard markers or reduced-representation approaches (Ajithkumar et al., 2024; Hollenbeck & Johnston, 2018; Launey & Hedgecock, 2001). This highlights the importance of carefully designed arrays for accurate, genome-wide genotyping in both *Ruditapes* species. Overall, the uniform MAF profiles, minimal missing data, and genome-wide marker coverage confirm the robustness of the array and its utility as a reliable genotyping platform across both species. Compared with genome-wide approaches such as GBS or low-pass sequencing, the SNP array provides a cost-effective and highly reproducible platform, minimizing missing data and facilitating standardized genotyping across populations and breeding programs (Darrier et al., 2019; Zenger et al., 2018).

**Figure 4:**
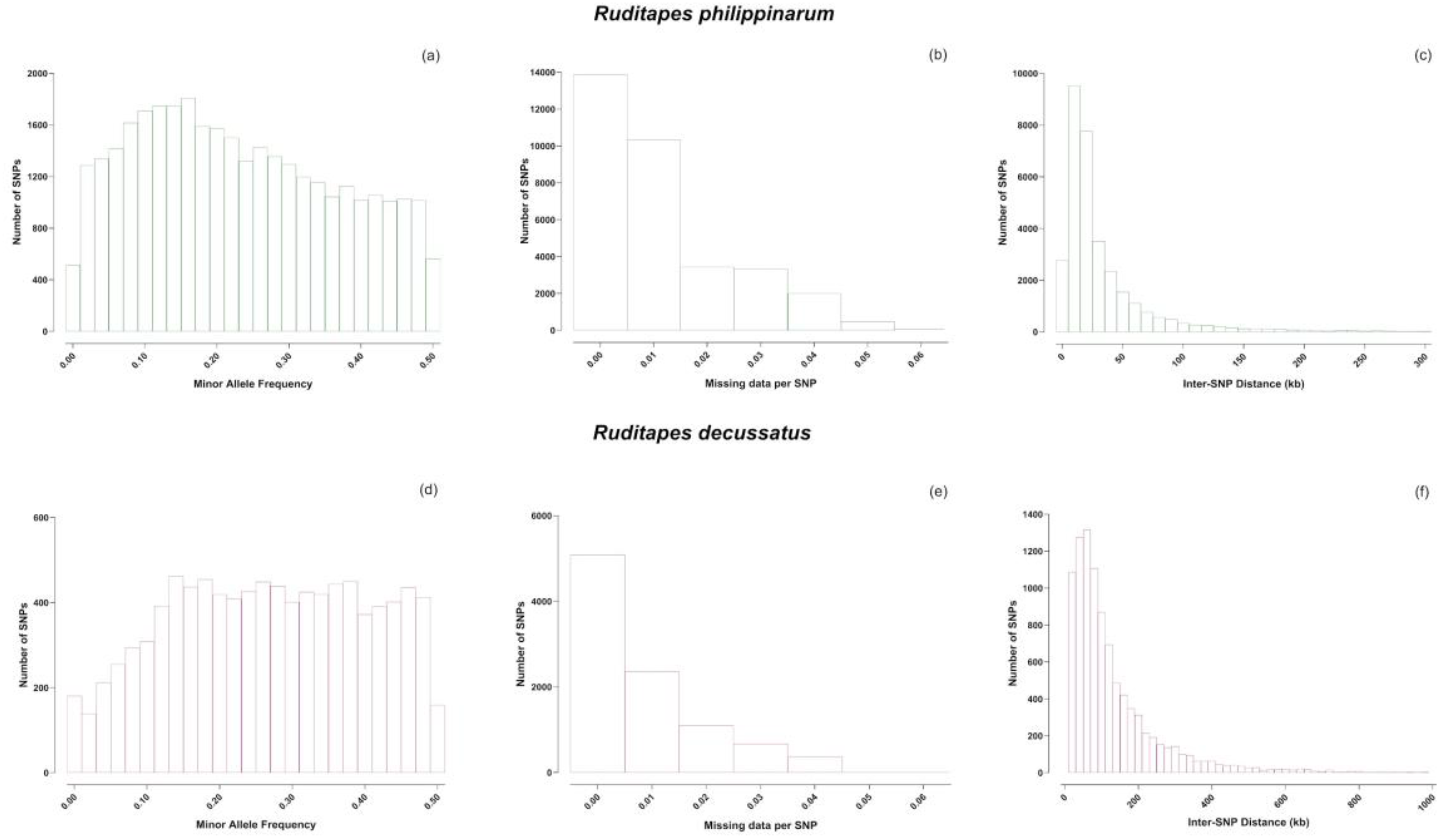
Quality parameters of the SNP Array for *R. philippinarum* (upper panel) and *R. decussatus* (lower panel). The distribution of minor allele frequencies (MAF > 0.05) is shown in (a) and (d); the proportion of missing data per SNP (≤ 5%) is shown in (b) and (e), and the distribution of inter-SNP distances is shown in (c) and (f).

### Genetic Differentiation Among Clam Populations

A comprehensive population genetics analysis of all genotyped populations is beyond the scope of this study. However, to assess the discriminatory power of the SNP chip with samples from wild populations, a subset comprising four *R. philippinarum* and all three *R. decussatus* wild populations was selected to evaluate the effectiveness of the 63K multispecies array for genetic stock discrimination. Principal component analysis (PCA) revealed a significant differentiation among *R. philippinarum* populations from different geographic origins (i.e. FR, PG, UK, and CHPF) (Figures 1, 5A), despite some intermingling in the borders. The modest fraction of the total variability explained by the first two axes (7% and 5.8% for PC1 and PC2, respectively) aligns with the previous observation and is consistent with expectations for an exotic species transferred to Europe recently (Chiesa et al., 2017; Gosling, 2008). Despite that, individuals cluster into distinct groups that roughly recapitulate populations, demonstrating the chip ability to discriminate between different genetic backgrounds. In *R. decussatus*, the three populations (ALG, VEN, SARD) were consistently separated according to their geographic distribution (Figures 1, 5B), as previously shown by Sambade et al. (2025). ALG population clustered within PC2 and whilst spread along PC1 (14.5% of the variance) was distinguished from VEN and SARD, while VEN and SARD populations were distinguished along PC2 (6% of the variance). Interestingly, some individuals from SARD were included with VEN suggesting some mixing between the populations (Carugati et al., 2024; Cruz et al., 2020). Population genetics analyses in bivalves are often challenging due to high larval dispersal, frequent anthropogenic translocations, and hatchery-based production, which can mask natural patterns of genetic differentiation. The SNP array provides a standardized tool to overcome these challenges, allowing robust detection of population structure even in recently admixed or managed stocks (Grouzdev et al., 2024; Hollenbeck & Johnston, 2018). Together, these results show that the array discloses the genetic differentiation patterns in both species, confirming its robustness as a powerful tool for studying genetic structure in wild populations.

**Figure 5.**
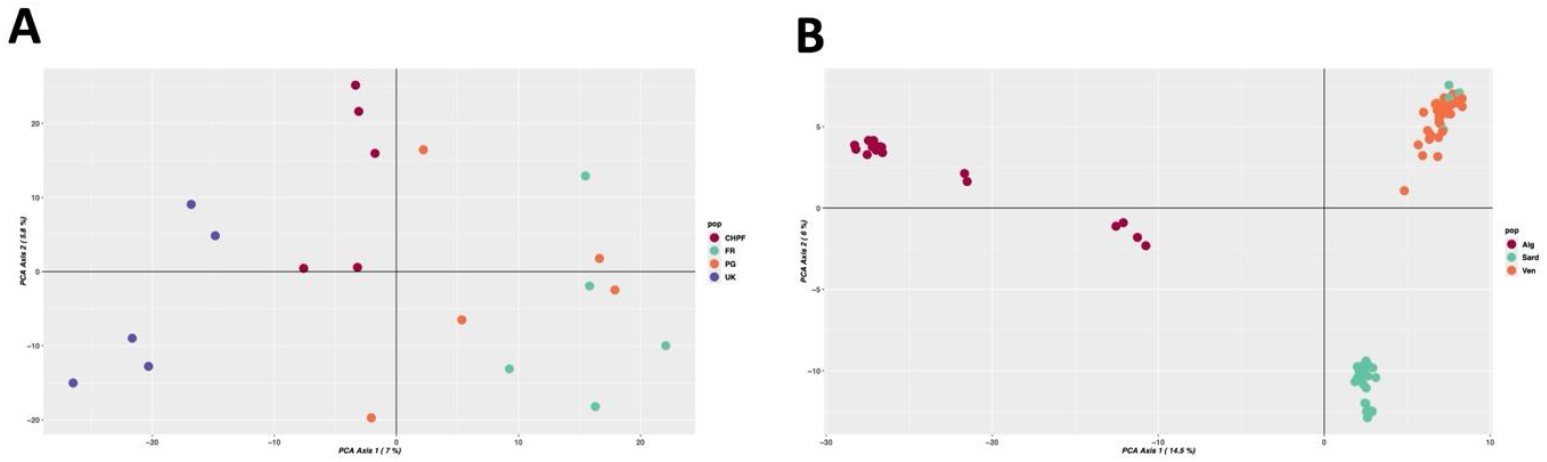
Principal component analysis (PCA) of selected populations of *R. philippinarum* (A) and *R. decussatus* (B). For panel A: CHPF, Chioggia, Italy; FR, Bay of Brest, France; PG, Ria de Aveiro, Portugal. For panel B: ALG, Algarve, Portugal; SARD, Sardinia, Italy; VEN, Venice, Italy.

### Validation of the SNP Array for Parentage Assignment in *R. philippinarum*

To further explore the applicability of the array, we estimated the genotyping error rate using a set of *R. philippinarum* families genotyped during the SNP array validation, for which family relationships were known. These samples were used to simulate different parentage scenarios and to evaluate the distribution of Mendelian mismatches as an indicator of genotyping accuracy and reliability of parental assignment. We evaluated parentage relationships using the full set of 33,453 SNPs for Manila clam with the Family Analysis Program (FAP), adapted to handle high-density SNP data, simulating three scenarios with increasing levels of uncertainty: *i)* true families; *ii)* one incorrect parent and *iii)* two incorrect parents. The analysis included 61 offspring from 36 full-sib families with known maternal and paternal genotypes. A clear and significant separation was observed among the three scenarios (Kruskal-Wallis test, H_(2)_ - p-value < 0.001, Figure 6 and Supplementary Table 1), indicating that the distribution of Mendelian mismatches is highly informative for assessing correct parental assignment. When both true parents were included, the proportion of Mendelian mismatches was low (average = 2.4%), with most offspring showing less than 3% mismatches (Figure 6, Suppl. Table 1). When only one true parent was present, the mismatch rate increased markedly to an average of 14.6% ranging between 13% and 16%. Finally, when both parents were randomly assigned, the mismatch rate reached 22% on average, ranging between 21% and 24%. These results demonstrate that the SNP array enables a clear distinction between correct and increasingly incorrect parental assignments based on Mendelian congruence, providing a robust framework for evaluating confidence in parentage assignment and estimate genotyping error rates in *R. philippinarum*. Overall, this array represents a valuable resource for applications requiring pedigree validation, such as the design and monitoring of controlled breeding programs and the development of traceable genetic lines in hatchery-based systems

**Figure 6:**
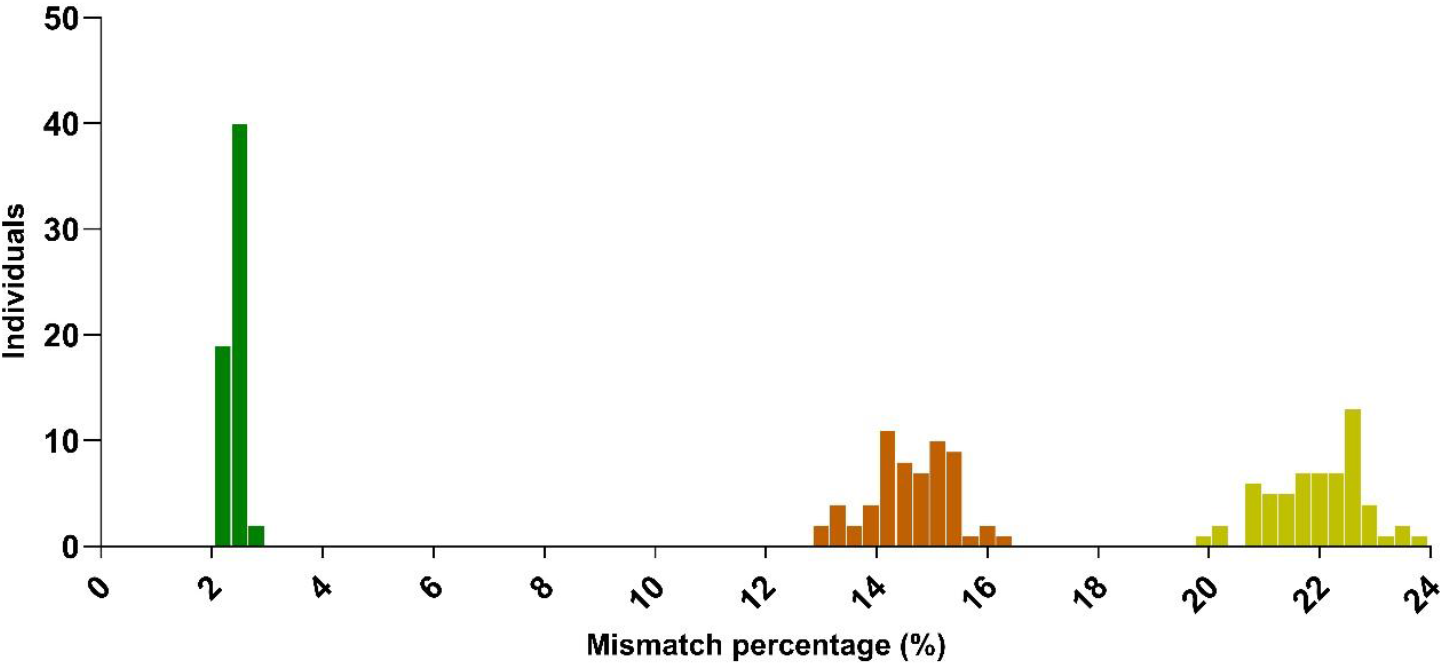
Distribution of genotype mismatch percentages across three simulated parentage assignment scenarios in *R. philippinarum*, based on 33,453 SNPs of the array. Green bars represent offspring with both parents correctly assigned, orange bars indicate offspring with only one true parent, and yellow bars correspond to cases where neither parent is correct.

## Conclusions

The development of a high-density SNP array for *R. philippinarum* and *R. decussatus* represents a major advancement for genomic applications in clam aquaculture. The array provides reliable and evenly distributed genomic coverage in both species, with high SNP conversion rates and strong genotyping quality, as demonstrated by MAF distribution, minimal missing data, population-level differentiation, and excellent parentage assignment accuracy. This performance confirms the robustness of the platform for multiple applications, including pedigree reconstruction, population genomics, stock discrimination and traceability. Beyond the validations presented here, the array will enable future GWAS, genomic selection, and studies on clam resilience to environmental and pathogenic stressors, supporting both breeding programs and conservation strategies for wild and cultured clam populations. By making this multispecies SNP array publicly available, we provide a long-needed genomic resource that will facilitate the development of genetically informed stock management and the sustainable evolution of clam aquaculture in Europe and beyond.

## FUNDING

This work was supported by the ‘‘Supporting Talent in ReSearch@University of Padua’’ grant for the project ASAP (funding from ‘‘Ministero dell’Istruzione e della Ricerca’’, CUP: C95F21009990001) awarded to LP; by the European Union grant IGNITION (GA 101084651) and the UK Research and Innovation (UKRI) awarded to MM, SFB and TB; by the European Union grant ShellFishBoost (GA: SBEP23_00051) awarded to LB, MM, LP, PM and SFB; this project was additionally funded under the National Recovery and Resilience Plan (NRRP), Mission 4 Component 2 Investment 1.4 - Call for tender no. 3138 of 16 December 2021, rectified by Decree n.3175 of 18 December 2021 of Italian Ministry of University and Research funded by the European Union –NextGenerationEU. SFB was supported through the CEEC Institutional program with ref. CEECINST /00027/2021/CP2789/CT0002; by the BBSRC Institute Strategic Programme grant (BBS/E/RL/230001A) awarded to TB;

## ACKNOWLEDGEMENTS

Authors are grateful to the hatchery SATMAR (France) and the hatchery Ecotapes Zeeland (Netherlands) for assistance in providing clam samples: in particular, authors are grateful to Mr Alexis Barreteau, Mr Jean-François Auvray and Ms Marina Martin for their continuous support over the past years.

## Notes

### Competing Interest Statement

The authors have declared no competing interest.

