## Supplementary material for "A dual-clam species 63K SNP array for sustainable production and conservation of wild resources": Figure S1; table S1

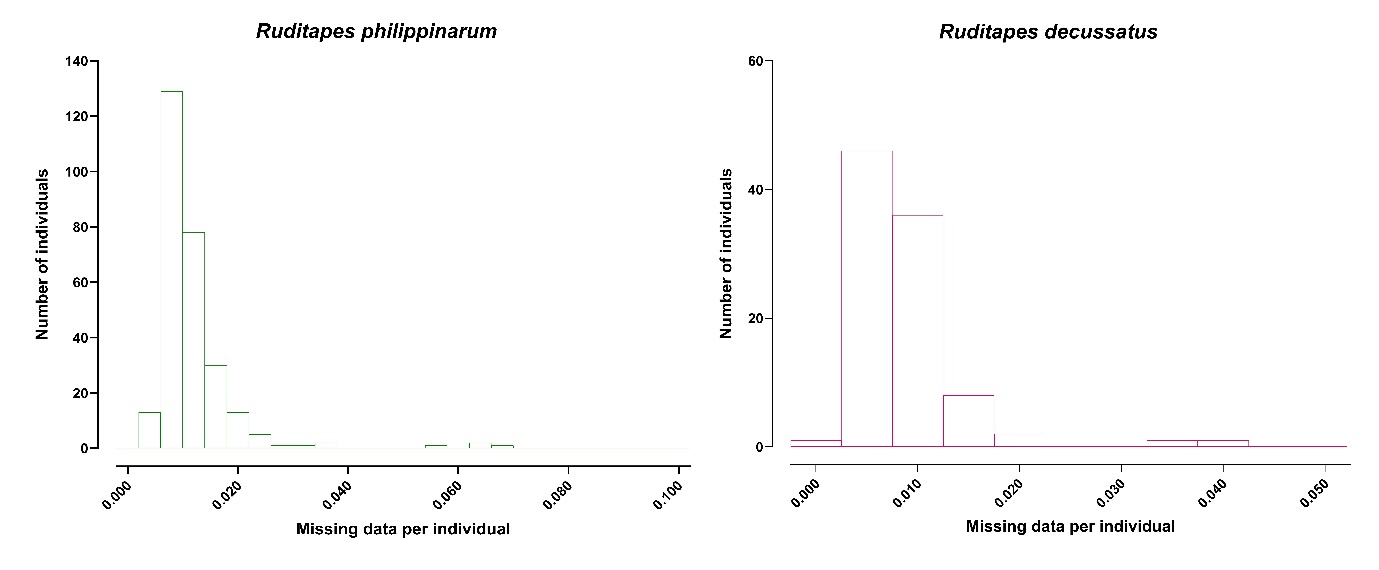

**Figure S1: Quality parameters of the SNP Array for *R. philippinarum* and *R. decussatus –*** Proportion of missing data per individuals

**Supplementary Table 1:** Statistical analysis on mismatch error across different parentage scenarios.

| *Kruskal-Wallis Test* | | | |
| --- | --- | --- | --- |
| Factor | Statistic | df | p |
| Parentage Scenarios | 963.2 | 2 | < .001 |

| *Dunn's Post Hoc Comparisons – Parentage Scenarios* | | | | | | | |
| --- | --- | --- | --- | --- | --- | --- | --- |
| Comparison | z | W_i_ | W_j_ | r_rb_ | p | p_bonf_ | p_holm_ |
| FS - HS | -6.891 | 31.00 | 359.5 | 1.000 | < .001 | < .001 | < .001 |
| FS - UR | -19.091 | 31.00 | 943.0 | 1.000 | < .001 | < .001 | < .001 |
| HS - UR | -28.097 | 359.50 | 943.0 | 1.000 | < .001 | < .001 | < .001 |
| *Note.*  Rank-biserial correlation based on individual Mann-Whitney tests. | | | | | | | |
